# *Drosophila* Filamin exhibits a mechano-protective role during nephrocyte injury via induction of hypertrophic growth

**DOI:** 10.1101/2021.08.20.457058

**Authors:** Sybille Koehler, Barry Denholm

**Affiliations:** Biomedical Sciences, University of Edinburgh, Edinburgh, Scotland, UK

## Abstract

Podocytes are highly specialized epithelial cells of the kidney glomerulus and are an essential part of the filtration barrier. Due to their position and function in the kidney, they are exposed to constant biomechanical forces such as shear stress and hydrostatic pressure. These forces increase during disease, resulting in podocyte injury and loss. The mechanism by which biomechanical forces are sensed and transduced to elicit an adaptive and protective response remains largely unknown. Here we show, using the *Drosophila* nephrocyte model, that the mechanosensor Cheerio (dFilamin) is central to this ‘mechano-protective’ mechanism. We found expression of an activated mechanosensitive variant of Cheerio induced hypertrophy and rescued filtration function in injured nephrocytes. Additional analysis with human Filamin B confirmed this mechano-protective role. We delineated the mechano-protective pathway downstream of Cheerio and found activation of TOR and Yorkie induce nephrocyte hypertrophy, whereas their repression reversed the Cheerio-mediated hypertrophy. Although Cheerio/Filamin B pathway mediates a mechano-protective role in the face of injury, we found excessive activity resulted in a pathological phenotype, indicating activity levels must be tightly controlled. Taken together, our data suggest that Cheerio acts via the TOR and YAP pathway to induce hypertrophic growth, as a mechano-protective response to nephrocyte injury.

## Introduction

Podocytes are highly specialized epithelial cells in the kidney glomerulus which together with fenestrated endothelial cells and the glomerular basement membrane (GBM) form the three-layered filtration barrier. The unique morphology of podocytes is exemplified by the formation of primary and secondary foot processes. These processes enwrap completely the glomerular capillaries and the adjacent foot processes of neighbouring podocytes interdigitate. This specialized arrangement facilitates the formation of a unique cell-cell-contact, the slit diaphragm (SD) (Arakawa, 1970; Pavenstädt et al., 2003). Interestingly, the glomerular filtrate has the highest extravascular flow rate in the human body and depends on the hydrostatic pressure difference across the filtration barrier (Kriz and Lemley, 2017). Podocyte morphology and attachment to the GBM is crucial to counteract these forces of flow and pressure.

Due to their position and function in the glomerular filter, podocytes face continual and fluctuating biomechanical forces of different types such as tensile forces induced by hydrostatic pressure in the capillaries and fluid shear stress in the filtration slit as well as in the Bowman’s space (Endlich et al., 2017). These biomechanical forces increase during diseases such as hypertension and diabetes, inflicting podocyte injury that result in changes to their morphology, detachment from the basement membrane and ultimately to their loss as they are shed into the primary urine. As post-mitotic cells, podocytes cannot be replenished, which leaves the capillaries blank and result in proteinuria, loss of protein from blood into urine. To avoid this, it is conceivable that podocytes possess mechanisms to monitor and adapt to fluctuations in biomechanical forces, which are up-regulated upon podocyte injury. Indeed, they contain a contractile actin-based cytoskeleton similar to that in smooth muscle cells suggesting their morphology is ‘plastic’ which might be an important component of the adaptive mechanisms required to withstand mechanical forces (Faul et al., 2007). This actin-based cytoskeleton plays a crucial role in the development and maintenance of podocyte morphology.

In general, the process of transducing biomechanical force into biochemical signal is called mechanotransduction (Endlich et al., 2017). Mechanosensors—as they detect biomechanical force—are essential for this process. Several classes of mechanosensors such as ion channels, proteins associated with the actin-cytoskeleton and proteins of the extracellular matrix, are already known (Endlich et al., 2017). In fact, a few proteins were already described to have a mechanosensor function in podocytes. Among them are the ion channels TRPC6 and P2×4 (Anderson et al., 2013; Forst et al., 2016) as well as the actin-associated proteins Cofilin, Paxillin and Filamin (Endlich et al., 2001; Greiten et al., 2021; Hayakawa et al., 2011; Okabe et al., 2021; Smith et al., 2014). However, detailed investigations into the functional roles of the mechanosensors in podocytes are scarce. In addition, the question whether additional proteins exhibit a mechanosensor function in podocytes remains unknown.

We and others have found the mechanosensor Filamin B to be upregulated upon podocyte injury in mice and in patient-derived glomerular tissue (Greiten et al., 2021; Koehler et al., 2020; Okabe et al., 2021). It is not known if this upregulation is a protective mechanism or induced as part of the pathological injury. Moreover, the functional role of Filamin B in podocytes remains largely unknown until today, as a global loss of Filamin causes embryonic (E14.5) or postnatal (directly after birth) lethality (Dalkilic et al., 2006; Feng et al., 2006; Hart et al., 2006; Zhou et al., 2010). Filamins are evolutionarily highly conserved, containing a N-terminal actin-binding domain and 24 β–sheet Ig domains, of which domain 16 to 24 contribute to the mechanosensor region (Razinia et al., 2012) **(Supp Figure 1A)**. Vertebrates have three Filamins (Filamin A, B and C), whereas the *Drosophila* and *C. elegans* genomes encode a single Filamin (Cheerio and FLN-1 respectively) (Razinia et al., 2012).

Here, we used the *Drosophila* nephrocyte model to unravel the functional role of Filamin B and to specifically address whether Filamin B has a mechano-protective role upon nephrocyte injury.

## Material and Methods

### Fly husbandry and generation

All flies were kept at 25°C for experiments. The nephrocyte-specific expression was achieved by mating UAS fly strains to the Sns-Gal4 strain.

Human Filamin B isoforms were amplified from a human embryonic kidney cDNA library using the following primers:

hFilamin B WT (full length):

fwd: 5’-CGCGGGACGCGTACCATGCCGGTAACCG -3’
rev: 5’-ATGCACGCGGCCGCTTAAGGCACTGTGAC– 3’
hFilamin B delta actin binding domain (ACB) (lacking bps 6 – 717):

fwd: 5’-CGCGGGACGCGTACC ATGCCGGCCAAGC-3’
rev : 5’-ATGCACGCGGCCGCTTAAGGCACTGTGAC– 3’
hFilamin B delta mechanosensor region (MSR) (truncation after 5445 bps):

fwd: 5’-CGCGGGACGCGTACCATGCCGGTAACCG -3’
rev: 5’-ATGCACGCGGCCGCTTATAGAACTGG – 3’
hFilamin B constructs were cloned into the pUAST vector with a C-terminal HA tag.

### Immunofluorescences of *Drosophila* tissue

*Drosophila* embryos were collected over 24 hrs and dechorionated in 50% bleach for 10 min. Following a washing step with H_2_O, they were fixed in 4% Formaldehyde/Heptane for 20 min. Heptane was then removed and replaced by Methanol, vortexed for 30 sec to devitellinized. The two phases separated after a minute and the embryos remained in the lower Formaldehyde phase. Both phases were removed and the embryos were washed with Methanol x3 for 20 min. The embryos were then washed with washing buffer (phosphate-buffered saline with 0.5 % BSA and 0.3 % Triton-X, PBS-Tx-BSA) x3 for 20 min, followed by overnight incubation at 4°C in primary antibody (see table 1). Primary antibody was removed from embryos and non-specific antibody binding was removed with 3x washes in PBS-Tx-BSA, the embryos were blocked in PBS-Tx-BSA + 5 % normal horse serum for 30 min, and then incubated with the appropriate secondary antibody for 1 h at room temperature (see table 1). Embryos were washed 3x 10 min in PBS-Tx-BSA before mounting in mounting medium (2.5% propyl gallate and 85% glycerol).

**Table 1:** List of antibodies.

| Name | Company/ Provider | Catalog nr. / Reference | Host Species | Dilution IF |
| --- | --- | --- | --- | --- |
| Cher | M. Uhlirova | (Külshammer and Uhlirova, 2013) | Rat | 1:100 |
| Duf | M. Ruiz-Gomez | (Weavers et al., 2009) | Rabbit | 1:100 |
| GFP | Abcam | ab6673 | Goat | 1:200 |
| HA tag | Proteintech | 51064-2-AP | Rabbit | 1:100 |
| Pyd | Developmental Studies Hybridoma Bank | PYD2 | Mouse | 1:25 |
| Albumin | Abcam | ab207327 | Rabbit | 1:25 |

**Table 2:** List of fly strains.

| **Fly strain** | **Origin** | **Purpose** | **Chromosome** |
| --- | --- | --- | --- |
| Cher-GFP-Trap | BDSC ID: 60261 | Expression pattern | 3. Chr. |
| UAS-Cher WT MSR | J. Ylänne (Huelsmann et al., 2016) | Overexpression | 3. Chr. |
| UAS-Cher inactive MSR (Cher closed) | J. Ylänne (Huelsmann et al., 2016) | Overexpression | 3. Chr. |
| UAS-Cher active MSR (Cher open) | J. Ylänne (Huelsmann et al., 2016) | Overexpression | 3. Chr. |
| UAS-Sns-RNAi | VDRC ID: 109442 | Knockdown | 2. Chr. |
| UAS-Duf-RNAi | VDRC ID: 27227 | Knockdown | 2. Chr. |
| Sns-Gal4; UAS-Dicer2 | Selfmade | Nephrocyte-specific driver | 2. + 3. Chr. |
| UAS-human Filamin B WT | Selfmade/ transgenics made by Genetivision | Overexpression | 3. Chr. |
| UAS- human Filamin B delta ACB | Selfmade/ transgenics made by Genetivision | Overexpression | 3. Chr. |
| UAS-human Filamin B delta MSR | Selfmade/ transgenics made by Genetivision | Overexpression | 3. Chr. |
| UAS-TOR-WT | BDSC ID: 7012 | Overexpression | 3. Chr. |
| UAS-TOR-DN | BDSC ID: 7013 | Overexpression | 2. Chr. |
| UAS-Rheb;UAS-TOR-WT<br>(TOR-OA) | BDSC ID:<br>80932 | Overexpression | 2. + 3. Chr. |
| UAS-arm.S10 (β-catenin<br>OA) | BDSC ID:4785 | Overexpression | X Chr. |
| UAS-pan.dTCFdeltaN (WNT<br>repressor) | BDSC ID:4784 | Overexpression | 2. Chr. |
| UAS-yki.S111A.S168A.S250A.V5<br>(YAP hyperactive) | BDSC ID:28817 | Overexpression | 3. Chr. |
| UAS-Hippo(dMST).Flag | BDSC ID:44254 | Overexpression | 3. Chr. |
| Oregon R |  | wildtype |  |
| W<sup>1118</sup> |  | wildtype |  |

Garland nephrocytes were prepared by dissecting a preparation containing the oesophagus and proventriculus (to which the garland nephrocytes are attached) from 3^rd^ instar larvae in haemolymph like buffer (HL3.1; 70 mM NaCl, 5 mM KCl, 1.5 mM CaCl_2_, 4 mM MgCl_2_, 10 mM NaHCO_3_, 115 mM Sucrose and 5 mM Hepes). The cells were fixed in 4 % Formaldehyde for 20 min, followed by three washing steps with PBS-Tx-BSA for 20 min. Incubation in primary antibody was at 4°C overnight. The next day, the preparation was washed x3 for 20 min, blocked with 5 % normal horse serum (in PBS-Tx-BSA) for 30 min, followed by incubation in the appropriate secondary antibody for 1 h at room temperature. The preparation was washed x3 for 20 min, before mounting in mounting medium.

Imaging was carried out using either a Zeiss LSM 800 confocal or a Leica SP8 confocal. Images were further processed using ImageJ (version 1.53c).

### FITC-Albumin assay

FITC-Albumin uptake assays were performed as previously published (Hermle et al., 2017; Koehler et al., 2020). Garland nephrocytes were prepared as described above and incubated with FITC-Albumin (0.2 mg/ml; Sigma-Aldrich, St. Louis, USA) for 1 min, followed by 20 min fixation with 4 % Formaldehyde. Nephrocytes isolated from Cheerio overexpression flies were used for additional antibody staining against Albumin (Cy3-labelled secondary antibody), as they express GFP. Antibody based staining was performed as previously described. For comparative analysis, the exposure time of experiments was kept identical. The mean intensity of FITC-Albumin or the Cy3-labelled FITC-Albumin was measured and normalized to control cells. Images were also used to quantify nephrocyte size by measuring the area of each cell.

### AgNO_3_ toxin assay

Flies of the appropriate genotype were allowed to lay eggs for 24 hrs at 25°C on juice plates with yeast. Plates with eggs were incubated at 18°C for 24 hrs. A defined number of 1^st^ instar larvae (usually 20) were then transferred to juice plates supplemented with yeast paste containing AgNO_3_ (2 g yeast in 3.5 ml 0.003% AgNO_3_). Plates were kept at 25°C for 4 days to let larvae develop and pupate. The number of pupae was counted daily between days 5-9 after egg laying and percentage pupation (relative to the number of larvae added to plate) was calculated.

### Statistical analysis

To determine statistical significance Graph Pad Prism software version 8 for Mac (GraphPad Software, San Diego, CA) was used. All results are expressed as means ± SEM. Comparison of more than two groups with one independent variable was carried out using one-way ANOVA followed by Tukey’s multiple comparisons test. A *P-value* < 0.05 was considered statistically significant.

## Results

### Cheerio is expressed in nephrocytes and co-localizes with the nephrocyte diaphragm proteins Duf and Pyd

Previous studies showed Filamin B levels to be elevated in patients suffering from focal segmental glomerulosclerosis (FSGS) and in injured murine podocytes, but its functional role and whether the increased expression levels are protective or pathogenic, remain elusive (Koehler et al., 2020; Okabe et al., 2021). Hence, within this study, we investigated the functional role of the *Drosophila* Filamin homologue, Cheerio, in more detail. To do so, we used the nephrocyte model, which shows a high morphological and functional similarity with mammalian podocytes and is widely used as a model for podocytes (Hermle et al., 2017; Koehler et al., 2020; Weavers et al., 2009).

We found that Cheerio is expressed from late embryonic stages and maintained throughout the life cycle in *Drosophila* garland nephrocytes. Interestingly, it partly co-localized with the nephrocyte diaphragm protein Dumbfounded (Duf/dNeph) suggesting it is present at the nephrocyte diaphragm and might be part of the nephrocyte diaphragm multi-protein complex **(Figure 1A,B)**. Based on its expression in nephrocytes and subcellular localization to the nephrocyte diaphragm we reasoned that the nephrocyte would be an excellent system to model Cheerio/Filamin B function.

### Elevated active Cheerio levels resulted in nephrocyte hypertrophy

β1-Integrin, a transmembrane protein that forms part of the extracellular matrix, is crucial for mechanotransduction. Both Filamin and Cheerio are able to bind β1-Integrin and the actin cytoskeleton. Filamin/Cheerio therefore provide a functional link between the extracellular matrix and the actin-cytoskeleton and allow extracellular force to be transduced into the cell(Razinia et al., 2012). The interaction with ß1-Integrin is mediated by a force induce conformational change of Filamin, which results in access to domain 21 (Filamin) (Sutherland-Smith, 2011). To investigate the functional role of Cheerio and to mimic the elevated Filamin B levels observed upon podocyte injury, we assayed three different Cheerio variants in nephrocytes (Huelsmann et al., 2016) **(Supp. Figure 1A)**. Here, we focused on the functional role of the mechanosensor region (MSR). In addition to the wildtype MSR (Cher wildtype), we used a ‘closed’ variant which requires greater mechanical force to unmask the binding sites for downstream targets of Cheerio, between domains 15/16 and 17 /18, and is therefore less active (Cher Inactive) (Huelsmann et al., 2016; Nakamura et al., 2006). In addition, we used an ‘open’ variant which requires less mechanical force to unmask the interaction sites of downstream targets and is therefore more active (Cher Active). The mutations introduced into Cheerio to generate the active variant were reported to cause an enhanced integrin binding (Lad et al., 2007).

To unravel the functional role of Cheerio-MSR, we expressed the Cheerio variants with a nephrocyte-specific driver (Sns-Gal4). We first analysed the subcellular expression pattern of each variant using the C-terminal GFP tag. Cher wildtype and Cher Inactive were localized throughout the nephrocyte cytoplasm. In contrast, Cher Active accumulated predominantly at the nephrocyte periphery, indicative of localization at the nephrocyte diaphragm **(Figure 1C)**.

Upon expression of the different Cheerio variants nephrocyte diaphragm integrity and morphology were not changed **(Figure 1D,E)**. In contrast, nephrocyte size increased significantly when expressing Cher Active, but not Cher wildtype or Cher Inactive **(Figure 1F)**. We also analysed nephrocyte function. The normal function of nephrocytes is to remove toxins from the haemolymph by filtration and endocytosis; processes which require an intact and functional nephrocyte diaphragm (Weavers et al., 2009). These processes can be assayed by (i) measuring FITC-Albumin uptake in isolated nephrocytes and (ii) monitoring the ability of nephrocytes to clear ingested AgNO_3_ (a toxin) from the haemolymph in the intact animal (haemolymph borne AgNO_3_ reduces viability and slows development, therefore haemolymph AgNO_3_ concentration can be assessed indirectly by its impact on these parameters). We used both assays to determine the effects of Cheerio variants on nephrocyte function **(Figure 1G,H)**. No defects were found for any of the variants in the FITC-Albumin uptake assay **(Figure 1G, Supp. Figure 1B)**. In line with this, we could not detect any significant delay in pupation behaviour for the different Cheerio variants in comparison to control flies **(Figure 1H)**. When we compared the pupation behaviour of the different genotypes fed on AgNO_3_ with animals fed on normal yeast we also could not detect any differences **(Figure 1I)**. In summary, the expression of active Cheerio did not result in a pathological phenotype (i.e. there was no impact on nephrocyte morphology or filtration function), but stimulated growth of the nephrocyte. This, along with the fact that hypertrophic growth is a protective mechanism during podocyte injury (Kriz and Lemley, 2015), suggest increased Cheerio/Filamin activity levels is a protective rather than pathogenic response.

**Figure 1:**
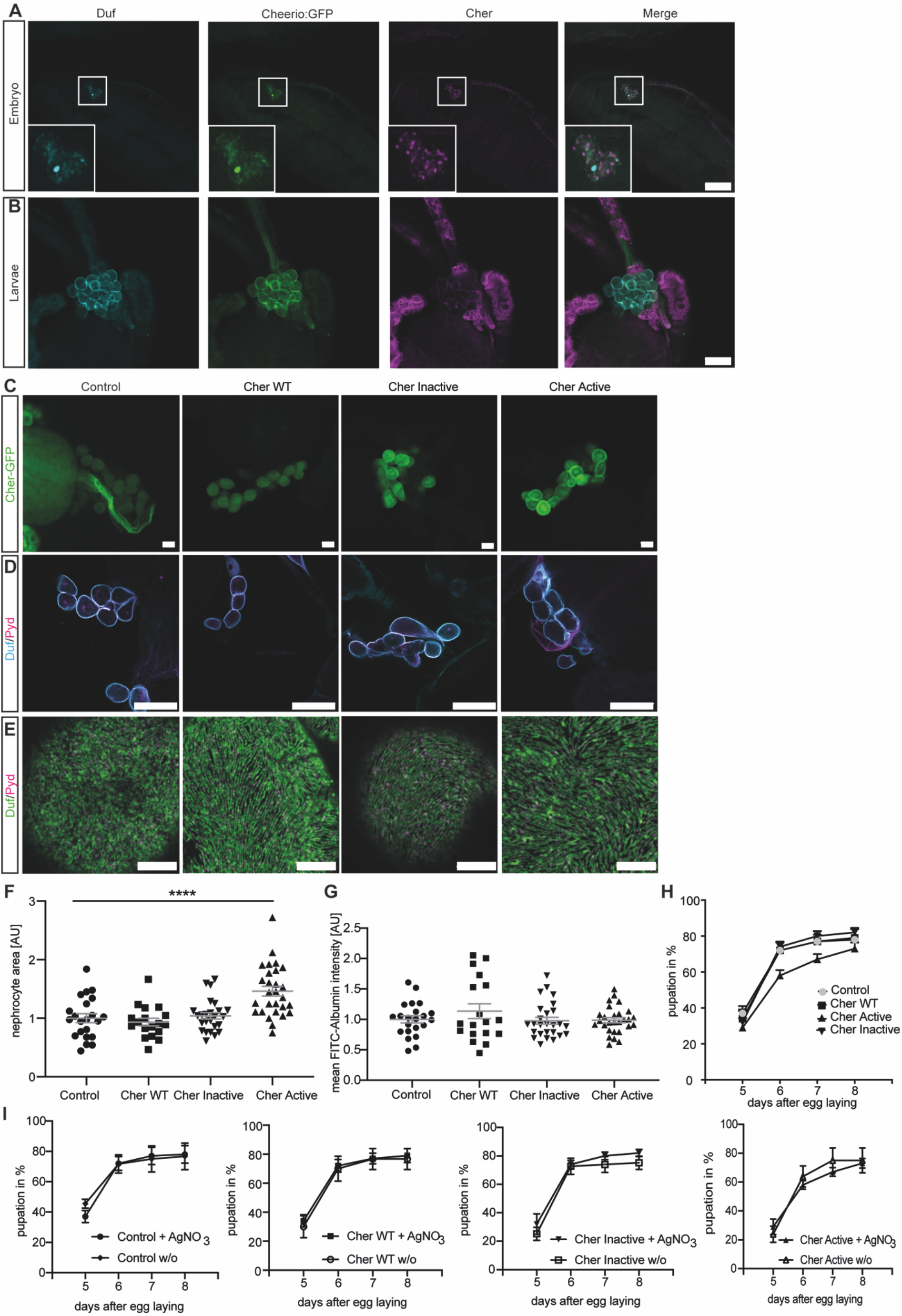
Elevated Cheerio-MSR levels result in nephrocyte hypertrophy. **A**,**B** Immunofluorescence staining confirmed the expression of Cheerio (green: Cheerio:GFP; magenta: anti-Cheerio antibody) in *Drosophila* wildtype embryos (**A**) and garland nephrocytes isolated from 3^rd^ instar larvae (**B**). Co-staining with a Duf antibody (cyan) revealed partial co-localisation with Cheerio. Scale bar = 50 µm (**A**) and 25 µm (**B**). **C** Cheerio mutants were expressed specifically in nephrocytes and are fused to a C-terminal GFP. Confocal microscopy revealed a cytoplasmic localization of wildtype (WT) and inactive Cheerio (Cher), while the active variant shows an accumulation at the cell cortex. Scale bar = 25 µm. Control: w;*sns*-Gal4;UAS-*dicer2*, Cher wildtype: w;*sns*-Gal4/UAS-*cher-wildtype-MSR*;UAS-*dicer2*/+, Cher Inactive: w;*sns*-Gal4/+;UAS-*dicer2*/UAS-*cher-inactive-MSR*, Cher Active: w;*sns*-Gal4/+;UAS-*dicer2*/UAS-*cher-active-MSR*. **D** Duf and Pyd stainings did not show any changes in morphology after expressing the Cheerio variants. Scale bar = 25 µm **E** Also, high resolution imaging using Airyscan and confocal microscopy did not reveal any differences when compared to control cells (w;*sns*-Gal4;UAS-*dicer2*). Scale bar = 5 µm. **F** Measuring nephrocyte size resulted in a significant size increase in nephrocytes expressing the active Cheerio variant relative to control cells. One-way ANOVA plus Tukey’s multiple comparisons test: ****: p < 0.001. **G** FITC-Albumin uptake assays did not reveal any differences. **H, I** The AgNO_3_ toxin assay showed a slightly delayed pupation in flies expressing the active Cheerio variant in nephrocytes, however this difference was not significant.

**Supp. Figure 1:**
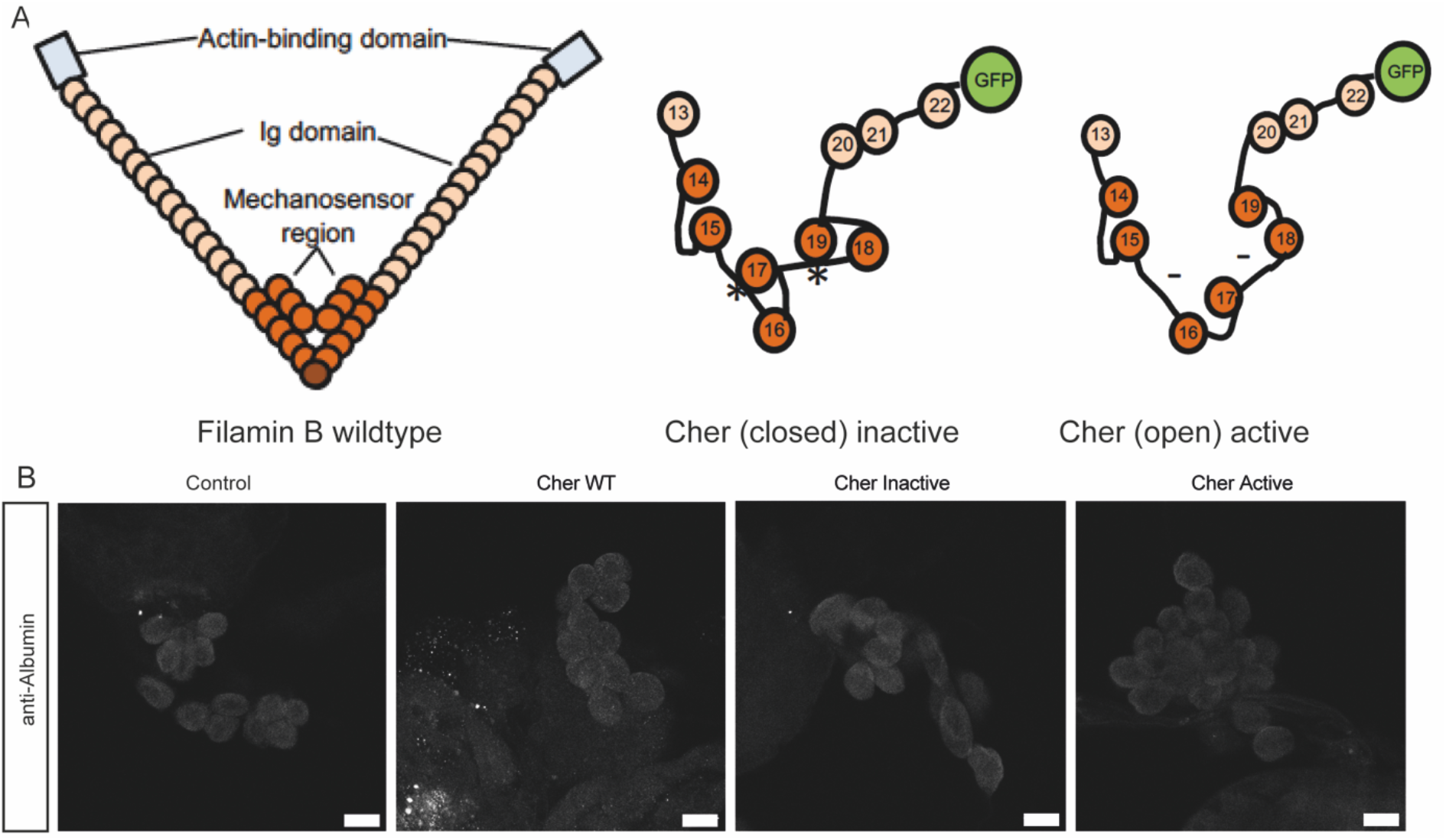
Cheerio variants. **A** Schematics showing the human Filamin B wildtype with its actin-binding domain (light blue), the IgG domains (light orange) and the mechanosensor region (orange). The Cheerio variants used within this study were generated by replacing the sequences between domain 15 and 16 as well as 17 and 18 (represented by asterisk and hyphen). Modified from Huelsmann et al. (Huelsmann et al., 2016) **B** FITC-Albumin assays did not reveal any differences between control and Cheerio expressing nephrocytes. Due to the GFP-tag FITC-Albumin was visualized by using an Albumin antibody. Scale bar = 25 µm. Control: w;*sns*-Gal4;UAS-*dicer2*, Cher wildtype: w;*sns*-Gal4/UAS-*cher-wildtype-MSR*;UAS-*dicer2*/+, Cher Inactive: w;*sns*-Gal4/+;UAS-*dicer2*/UAS-*cher-inactive-MSR*, Cher Active: w;*sns*-Gal4/+;UAS-*dicer2*/UAS-*cher-active-MSR*.

### Cheerio exhibits a protective role during nephrocyte injury

We next investigated whether Cheerio exhibits a similar protective role upon nephrocyte injury. Loss of either of the nephrocyte diaphragm components Sns (dNephrin) or Duf resulted in a severe nephrocyte injury that includes both morphological and functional disturbances (Weavers et al., 2009; Zhuang et al., 2009). In detail, nephrocyte diaphragm integrity was severely disrupted and cell size reduced, while FITC-Albumin uptake is significantly decreased and the onset of pupal development is severely delayed in the AgNO_3_ toxin uptake assay. In line with our hypothesis that Cheerio provides a protective role, we found that endogenous Cheerio translocated from the cytoplasm to the nephrocyte cortex upon Sns or Duf depletion **(Figure 2A, arrowheads)**.

To assess if Cheerio has a beneficial effect during injury, we induced injury (by depleting Sns or Duf) whilst simultaneously expressing the active or inactive Cheerio-MSR variants **(Figure 2, Supp Figure 2,3)**. Filtration function was restored as shown with the FITC Albumin uptake assays **(Figure 2D, Supp. Figure 3B)**, but nephrocyte morphology and size could not be rescued by expressing the active Cheerio **(Figure 3B,C)**. Active Cheerio also rescued the delayed pupation behaviour observed when depleting Duf **(Figure 2E,F)**. Expression of inactive Cheerio only resulted in a rescue of the delayed pupation in nephrocytes lacking Duf **(Supp. Figure 2A,B,C,D,E, Supp. Figure 3A)**.

**Figure 2:**
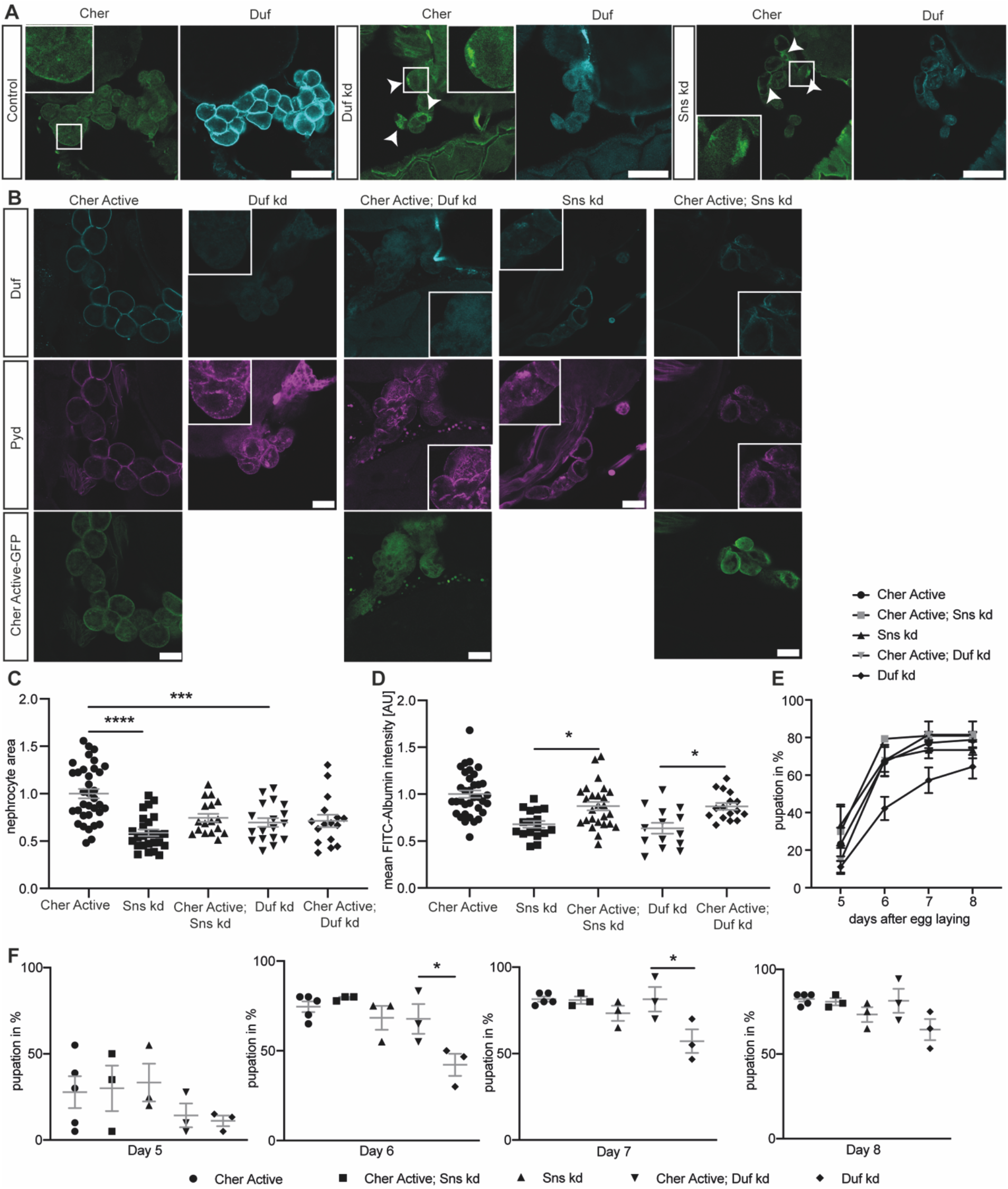
Active cheerio partially rescues the nephrocyte phenotype during injury. **A** Cheerio localisation was examined in injured nephrocytes. By depleting Duf and Sns, nephrocytes present with a severe morphological phenotype, as Duf expression and localisation is severely disrupted (cyan). Comparing Cheerio expression and localisation to control cells revealed an accumulation of Cheerio at the cell cortex in injured cells. Scale bar = 50 µm. **B** Active Cheerio was combined with Duf or Sns RNAi and showed no rescue of the nephrocyte morphology as depicted by Duf (cyan) and Pyd (magenta) staining. Scale bar = 25 µm. Cher active: w;*sns*-Gal4/+;UAS-*cher-active-MSR*/+, Duf kd: w;*sns*-Gal4/UAS-*duf*-RNAi;UAS-*dicer2*/+; Cher active; Duf kd: w;*sns*-Gal4/UAS-*duf*-RNAi;UAS-*cher-active-MSR*/+, Sns kd: w;*sns*-Gal4/UAS-*sns*-RNAi;UAS-*dicer2*/+; Cher active; Sns kd: w;*sns*-Gal4/UAS-*sns*-RNAi;UAS-*cher-active-MSR*/+. **C** Nephrocyte size could not be rescued by simultaneous expression of active Cheerio and Duf or Sns RNAi. Cher Active was used as control and all other genotypes are depicted as relative values of Cher Active. **D** FITC-Albumin uptake was restored in both, Duf and Sns depleted nephrocytes after expression of active Cheerio. One-way ANOVA plus Tukey’s multiple comparisons test: ***: p < 0.001. **E**,**F** The AgNO3 toxin assay revealed a rescue of the delayed pupation by expressing active Cheerio in combination with Duf or Sns RNAi. One-way ANOVA plus Tukey’s multiple comparisons test: *: p < 0.05.

**Supp. Figure 2:**
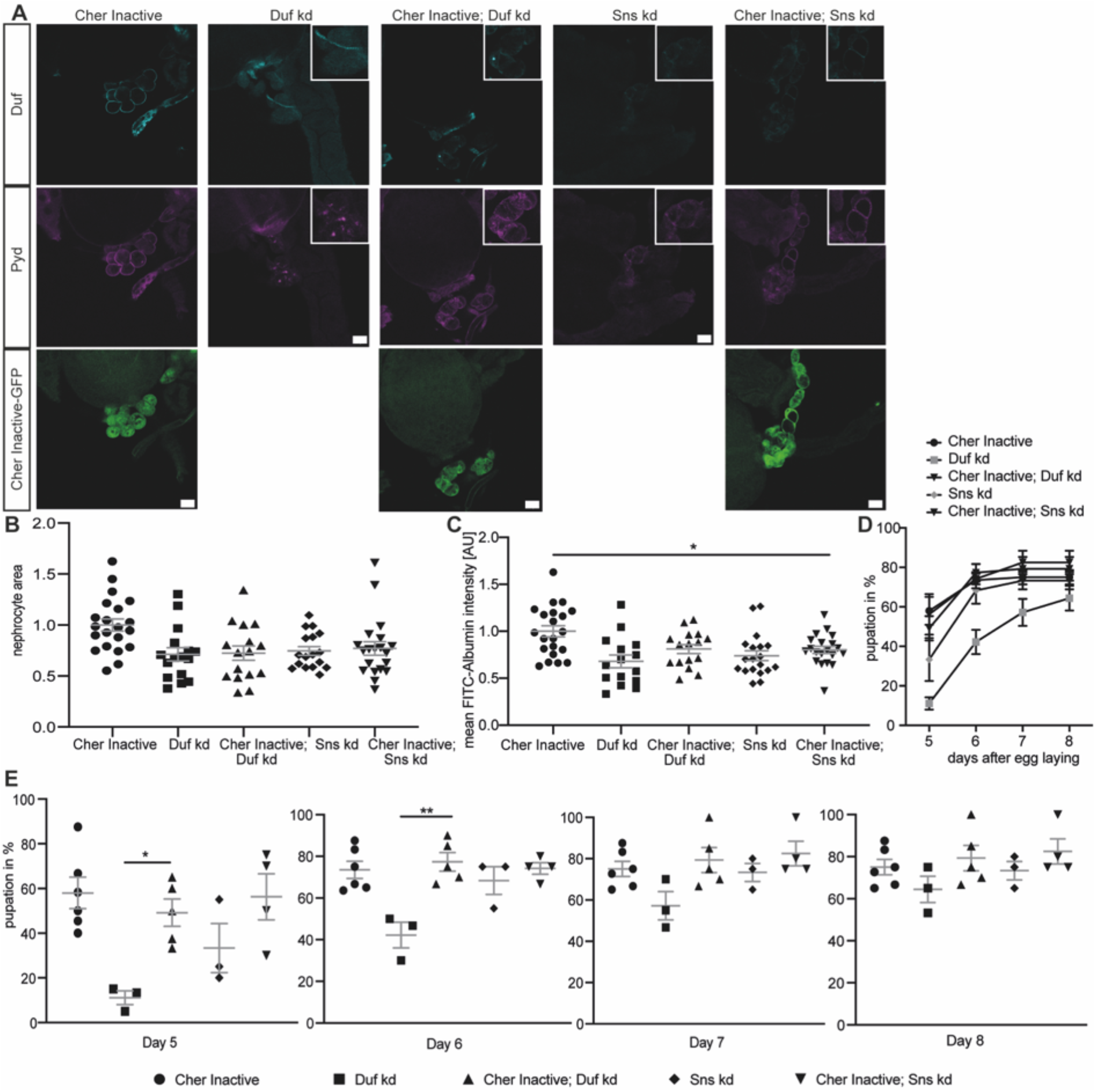
Inactive Cheerio partially restores filtration function during injury. **A** Inactive Cheerio MSR was combined with Duf or Sns RNAi and revealed no restoration of the nephrocyte diaphragm integrity, when compared to Duf or Sns knockdown. Scale bar = 25 µm. Cher inactive: w;*sns*-Gal4/+;UAS-*cher-inactive-MSR*/+, Duf kd: w;*sns*-Gal4/UAS-*duf*-RNAi;UAS-*dicer2*/+; Cher inactive; Duf kd: w;*sns*-Gal4/UAS-*duf*-RNAi;UAS-*cher-inactive-MSR*/+, Sns kd: w;*sns*-Gal4/UAS-*sns*-RNAi;UAS-*dicer2*/+; Cher inactive; Sns kd: w;*sns*-Gal4/UAS-*sns*-RNAi;UAS-*cher-inactive-MSR*/+. **B** Measuring nephrocyte size did not reveal a rescue in cells expressing the inactive Cheerio together with the Sns and Duf RNAi. Cher Inactive was used as control and all genotypes are depicted relative to Cher Inactive. **C** FITC-Albumin assays also did not show a rescue for both genotypes, inactive Cheerio combined with Duf or Sns RNAi. One-way ANOVA plus Tukey’s multiple comparisons test: *: p < 0.05. **D**,**E** The AgNO_3_ toxin assay however showed a significant rescue of the delayed pupation by expressing the inactive Cheerio MSR in a Duf knockdown background. One-way ANOVA plus Tukey’s multiple comparisons test: *: p < 0.05; **: p < 0.01.

**Figure 3:**
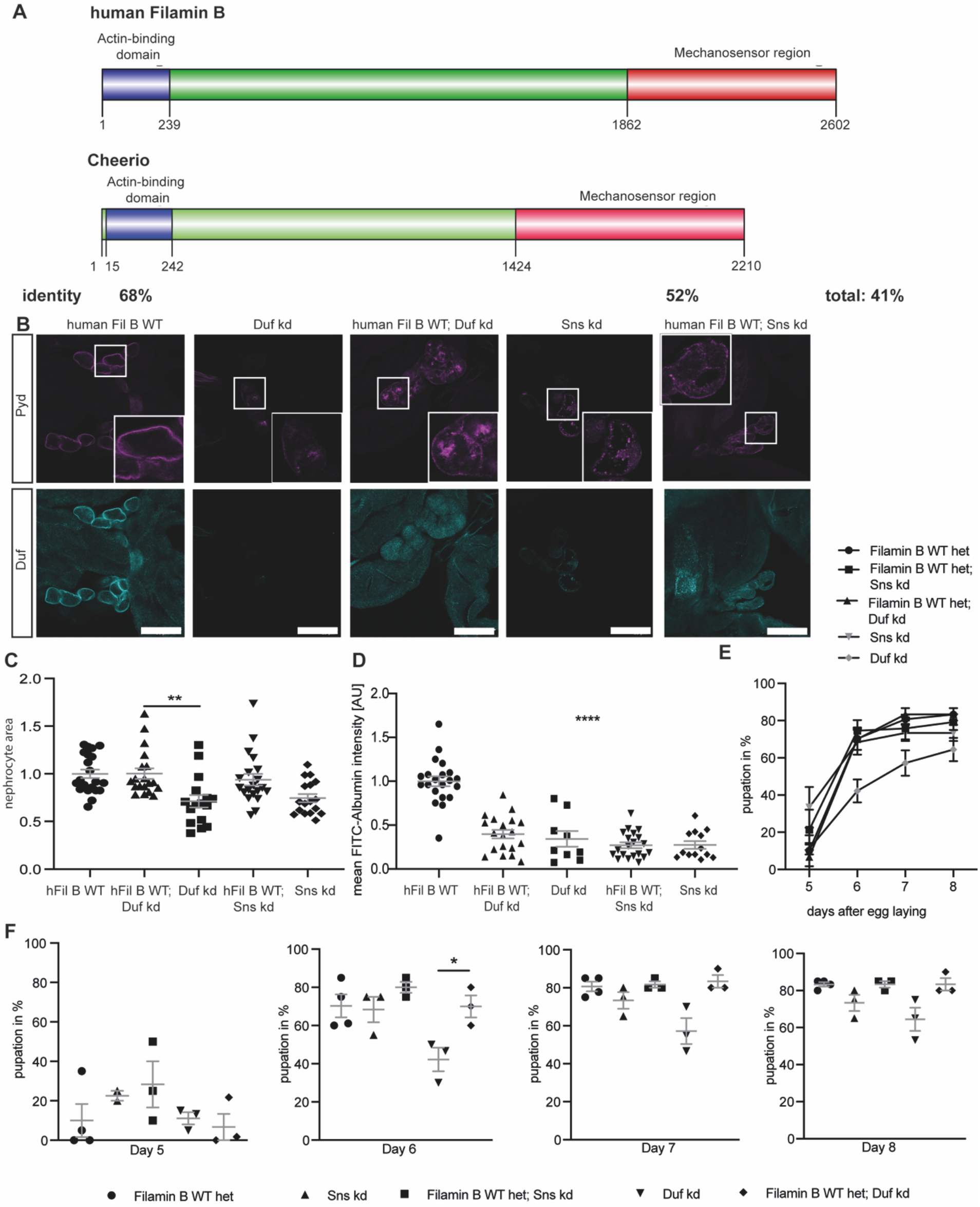
Human Filamin B rescues nephrocyte size and function. **A** Schematic of human Filamin B wildtype and *Drosophila* Cheerio wildtype. Both possess an actin-binding domain at the N-terminus, which shows a sequence identity of 68%. This domain is followed by the IgG domains, which form the mechanosensor region at the C-terminus. The MSR shows a sequence identity of 52%. **B** Expression of human Filamin B wildtype (human Fil B WT) in Duf and Sns depleted nephrocytes did not restore morphology. Scale bar = 25 µm. hFil B WT: w;*sns*-Gal4/+;UAS-*hFilamin B WT*/+; Duf kd: w;*sns*-Gal4/+;UAS-*duf-*RNAi/UAS-*dicer2*; hFil B WT; Duf kd: w;*sns*-Gal4/+;UAS-*hFilamin B WT*/UAS-*duf*-RNAi; Sns kd: w;*sns*-Gal4/+;UAS-*sns-*RNAi/UAS-*dicer2*; hFil B WT; Sns kd: w;*sns*-Gal4/+;UAS-*hFilamin B WT*/UAS-*sns*-RNAi. **C** Nephrocyte size was restored by expressing human Filamin B wildtype. One-way ANOVA plus Tukey’s multiple comparisons test: **: p < 0.01. **D** FITC Albumin uptake is not rescued by expression of the human Filamin B wildtype. One-way ANOVA plus Tukey’s multiple comparisons test: ****: p < 0.0001. **E** AgNO_3_ toxin assay revealed a rescue of the delayed pupation when human Filamin B wildtype was expressed simultaneously to the Duf or Sns RNAi. One-way ANOVA plus Tukey’s multiple comparisons test: *: p < 0.05.

**Supp. Figure 3:**
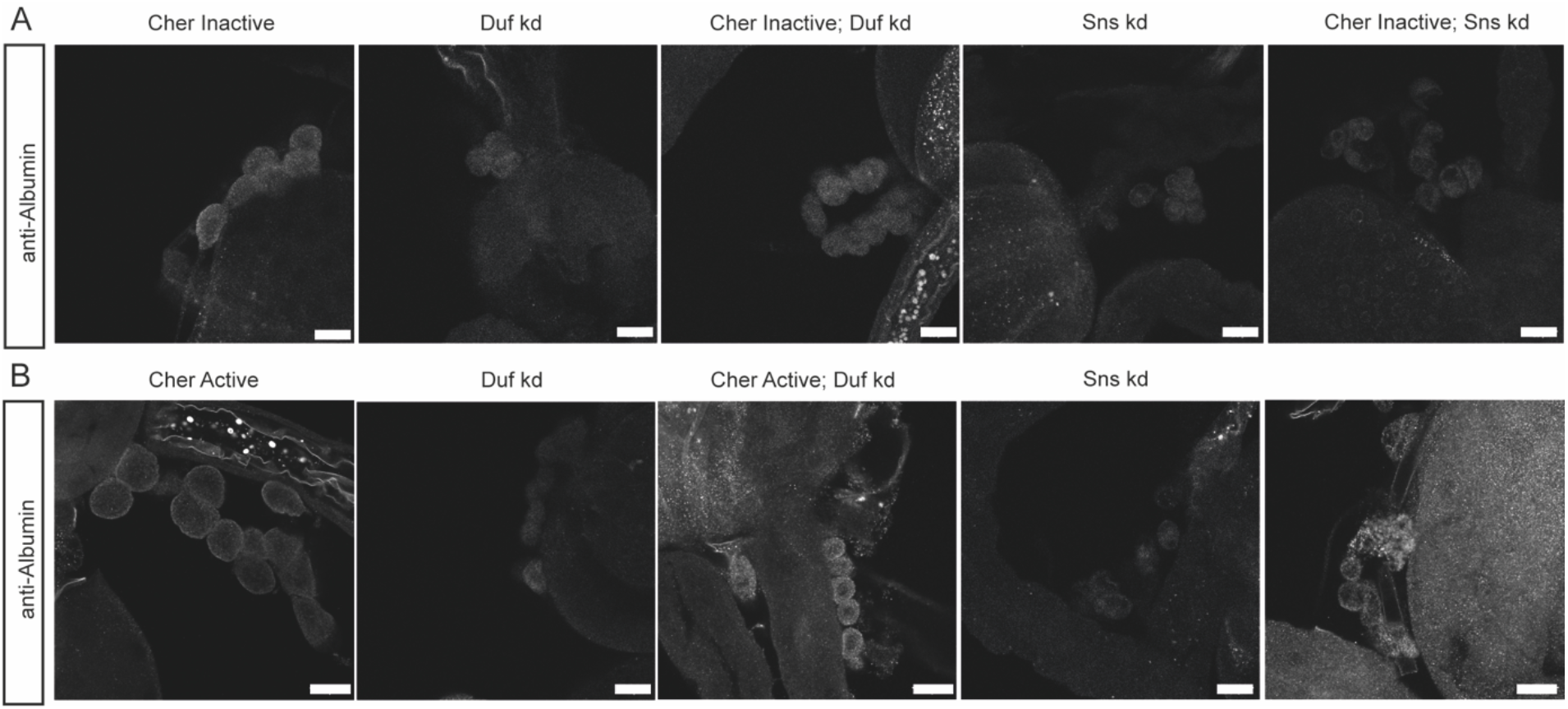
FITC-Albumin uptake assays of Cheerio rescue strains. **A** Expression of inactive Cheerio in a Duf or Sns knockdown background did not rescue the FITC-Albumin uptake. Scale bar = 25 µm. Cher inactive: w;*sns*-Gal4/+;UAS-*cher-inactive-MSR*/+, Duf kd: w;*sns*-Gal4/UAS-*duf*-RNAi;UAS-*dicer2*/+; Cher inactive; Duf kd: w;*sns*-Gal4/UAS-*duf*-RNAi;UAS-*cher-inactive-MSR*/+, Sns kd: w;*sns*-Gal4/UAS-*sns*-RNAi;UAS-*dicer2*/+; Cher inactive; Sns kd: w;*sns*-Gal4/UAS-*sns*-RNAi;UAS-*cher-inactive-MSR*/+. **B** Expression of active Cheerio in a Duf or Sns knockdown background partially restored the FITC-Albumin uptake. Scale bar = 25 µm. Cher active: w;*sns*-Gal4/+;UAS-*cher-active-MSR*/+, Duf kd: w;*sns*-Gal4/UAS-*duf*-RNAi;UAS-*dicer2*/+; Cher active; Duf kd: w;*sns*-Gal4/UAS-*duf*-RNAi;UAS-*cher-active-MSR*/+, Sns kd: w;*sns*-Gal4/UAS-*sns*-RNAi;UAS-*dicer2*/+; Cher active; Sns kd: w;*sns*-Gal4/UAS-*sns*-RNAi;UAS-*cher-active-MSR*/+.

The inactive variant translocated to and accumulated at the cell cortex during nephrocyte injury, suggesting an activation of Cheerio and its accumulation at the periphery during injury **(Supp Figure 2A)**. These data confirm a protective role of Cheerio-MSR during injury.

### Human Filamin B has a mechano-protective role during nephrocyte injury

The wildtype human Filamin B (hFilB WT) displays a high sequence identity with Cheerio within the actin-binding domain (ACB, 68%) and MSR (52%) **(Figure 3A)**. Hence, we generated flies expressing wildtype human Filamin B in nephrocytes exclusively. We first checked subcellular localization of Filamin B in nephrocytes and find it localized to the nephrocyte diaphragm **(Supp. Figure 4A)**. Interestingly, in contrast to Cheerio, expression of Filamin B did not result in hypertrophy, but caused a significantly increased FITC-Albumin uptake **(Supp. Figure 4A)**.

**Supp. Figure 4:**
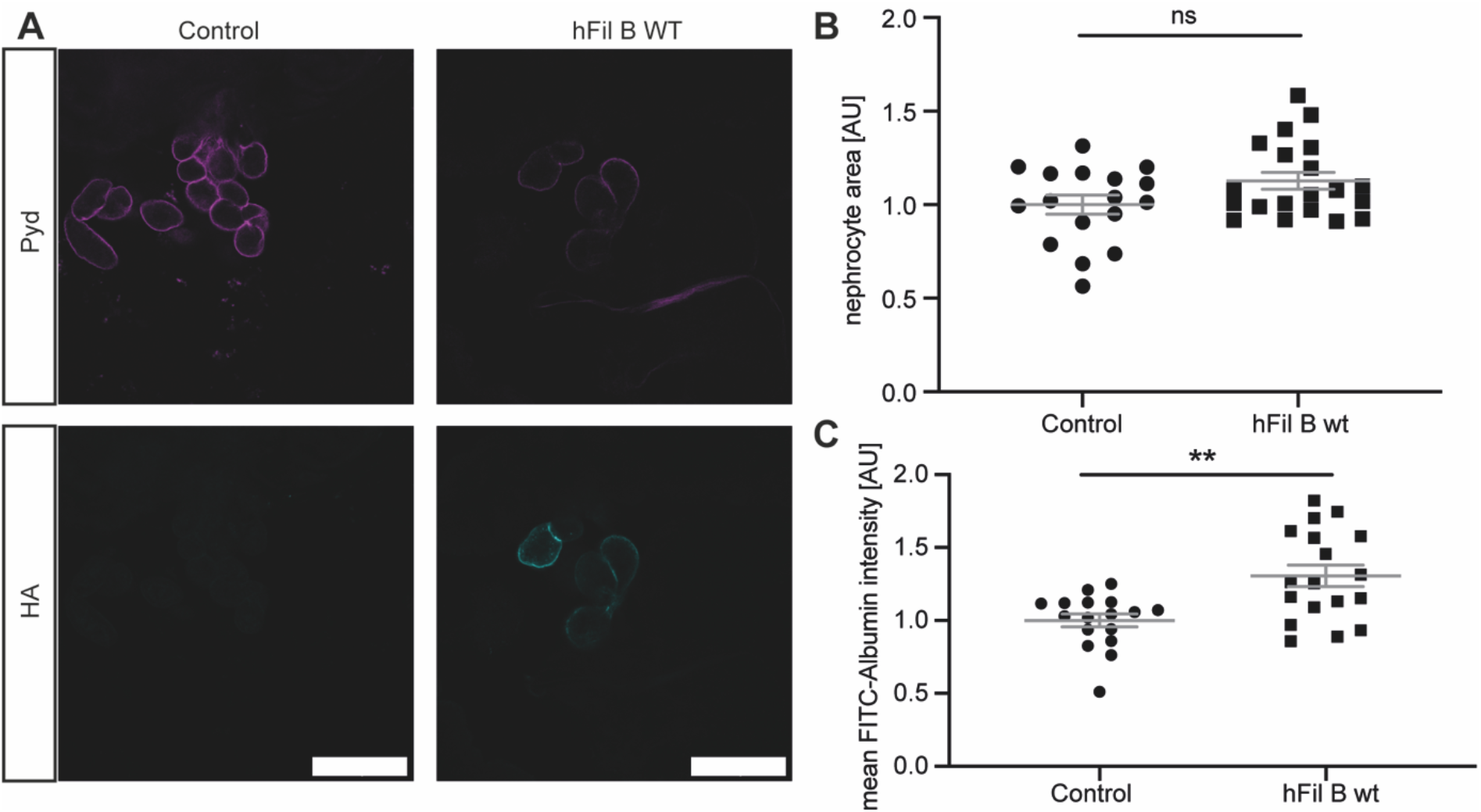
Human Filamin B localizes to the nephrocyte diaphragm and results in only a mild nephrocyte phenotype. **A** The human Filamin B construct expresses a C-terminal HA-tag, which was used for visualisation of the construct. Filamin B localized to the cell cortex. Scale bar = 25 µm. Control: w^1118^; h Fil B WT: w;*sns*-Gal4;UAS-*hFilamin B WT*. **B** Overexpression of human Filamin B wildtype resulted in a slight size increase, which was not significant. Control: w;*sns*-Gal4/+;MKRS/TM6B; h Fil B WT: w;*sns*-Gal4;UAS-*hFilamin B WT/+*. **C** FITC-Albumin assays reveal an increase of Albumin uptake in nephrocytes expressing human Filamin B. Student’s t-test: **: p < 0.01.

To investigate whether the protective role we have found for Cheerio is conserved for Filamin B, we generated transgenic flies in which Sns or Duf were depleted whilst simultaneously expressing human Filamin B.

We found a significant rescue of the nephrocyte size by simultaneous expression of human Filamin B wildtype, while morphology was partially restored **(Figure 3B,C)**. The onset of pupal development in the AgNO_3_ toxin assay was significantly reinstated in Duf depleted cells, validating a protective role of Filamin B during nephrocyte injury, although there was no rescue of the FITC Albumin uptake **(Figure 3E,F)**. In summary, neither Cheerio nor human Filamin B could restore morphology. The reduced cell size after Duf and Sns depletion could only be rescued by human Filamin B and not by Cheerio. Of note, the Cheerio constructs used do not possess the ACB, which could account for this functional difference. The functional assays revealed a significant rescue of the FITC-Albumin uptake phenotype when expressing Cheerio and a significant rescue of the delayed pupation behaviour when expressing either Cheerio or human Filamin B together with Duf depletion.

Taken together, these data indicate that human Filamin B exhibits a level of protection upon nephrocyte injury, although the phenotypes that are rescued are different to Cheerio.

### TOR and Yorkie signalling induce nephrocyte hypertrophy and appear to be downstream targets of Cheerio

The above results show a hypertrophy phenotype and protective role for Cheerio, while human Filamin B restored cell size and filtration function. To identify the downstream pathway that mediates these effects we performed a candidate screen examining three different pathways known to be involved in cell proliferation and tissue growth; TOR, Wingless (WNT) and Hippo (Huang et al., 2005; Laplante and Sabatini, 2012; Niehrs and Acebron, 2012). The over-activation of TOR, Yorkie (Yap/Taz; Hippo pathway) and armadillo (ß-Catenin; WNT pathway) in nephrocytes resulted in a significant size increase to a degree similar to active Cheerio, consistent with the notion that one or more of these pathways acts downstream of Cheerio **(Figure 4A,B,C)**. To help assess whether these pathways mediate the hypertrophic phenotype observed in nephrocytes expressing active Cheerio, we asked whether repression of these pathways is able to block the hypertrophic phenotype caused by active Cheerio. In detail, we expressed a dominant-negative TOR variant (TOR-DN), or overexpressed Hippo (dMST) to repress Yrk, in combination with expression of active Cheerio. (We also attempted to address whether WNT signalling was implicated by expressing a constitutive repressor version of the WNT effector pangolin (dTCF) (WNT DN). We were unable to interpret this experiment as WNT pathway repression resulted in the complete loss of nephrocytes, suggesting an important role of WNT signalling for the development and maintenance of nephrocytes, which is in line with previous reports for mammalian models (Carroll et al., 2005; Dai et al., 2009; Iglesias et al., 2007).) The repression of TOR signalling in active Cheerio expressing nephrocytes resulted in a significant size decrease back to normal levels (Cheerio inactive) **(Figure 4D)**. Similarly, repression of Yrk by expressing hippo in the presence of active Cheerio also resulted in a significant size decrease **(Figure 4E)**. These data are compatible with TOR and Yorkie acting in a pathway(s) downstream of Cheerio. However, TOR repression or Hippo overexpression in a wild-type background (i.e. without activated Cheerio), both resulted in a significant size decrease **(Figure 4F,G)**. Therefore, we cannot exclude the possibility that TOR and Yorkie pathways act independently of Cheerio.

**Figure 4:**
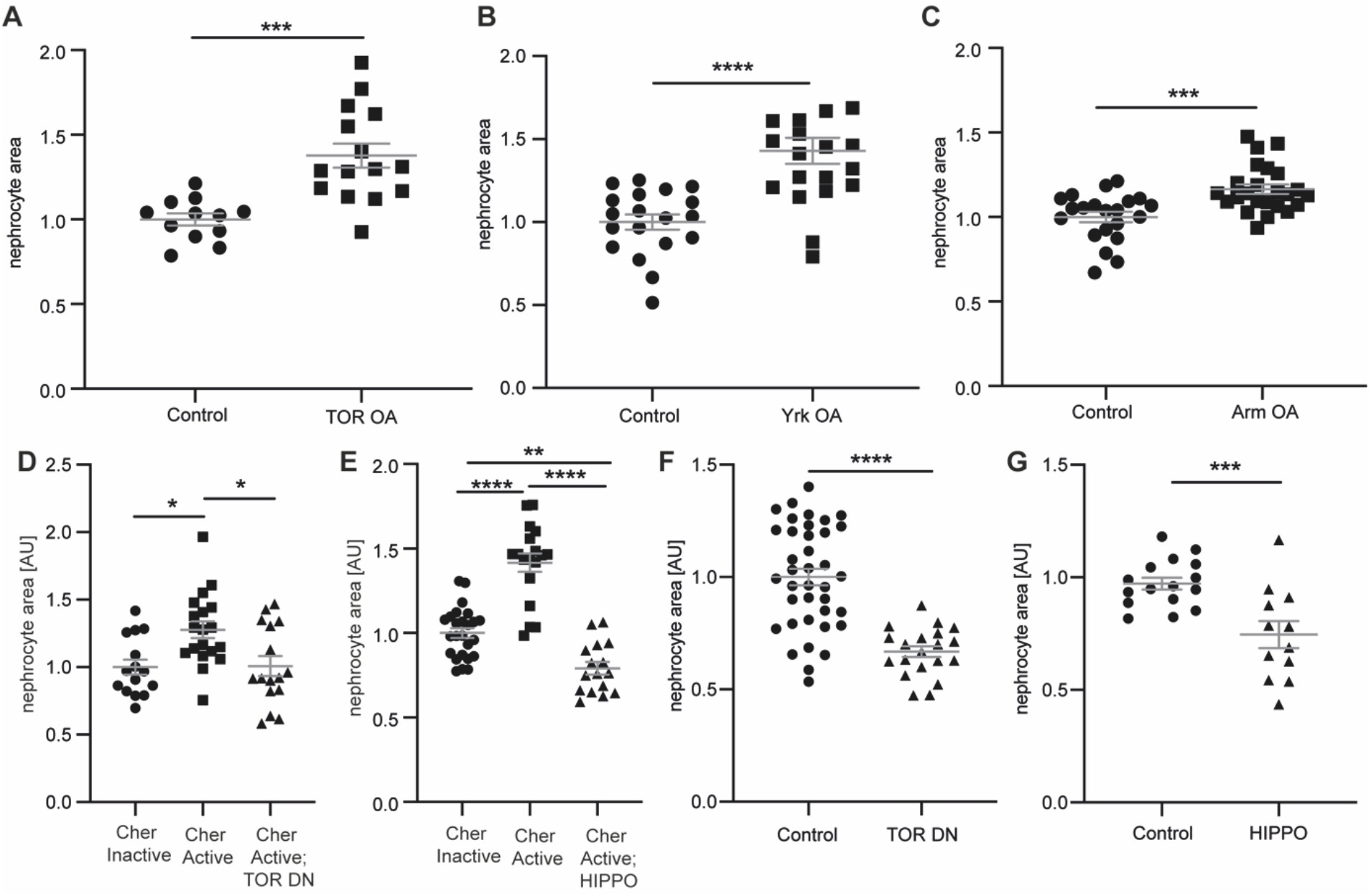
TOR and Hippo signalling mediate the hypertrophy phenotype in nephrocytes. **A** Over-activation of TOR signalling in nephrocytes caused a significant hypertrophy phenotype. Control: w;*sns*-Gal4/+;UAS-*dicer2*/+; TOR OA: w;*sns*-Gal4/UAS-*rheb*;UAS-*dicer2*/UAS-*tor-wildtype*. Student’s t-test: ***: p < 0.001. **B** Hyperactivation of Yorkie (Yap/Taz) in nephrocytes revealed a hypertrophy effect as well. Control: w;*sns*-Gal4/+;UAS-*dicer2*/+; Yrk OA: w;*sns*-Gal4/+;UAS-*dicer2*/UAS-*yki*.*S111A*.*S168A*.*S250A*.*V5*. Student’s t-test: ****: p < 0.0001. **C** Expression of constitutively active Armadillo (ß-Catenin) resulted in a significant size increase in nephrocytes. Control: w;*sns*-Gal4/+;UAS-*dicer2*/+; WNT OA: UAS-*arm*.*S10/*w;*sns*-Gal4/+;UAS-*dicer2*/+. Student’s t-test: ***: p < 0.001. **D** Quantification of the cell size based on GFP-tags to the Cheerio constructs revealed a significant size increase of Cher Active when compared to Cher Inactive cells. This increase was completely reversed when TOR signalling was inhibited with a dominant negative TOR variant (TOR DN). Cher Inactive: w;*sns*-Gal4/+;UAS-*cher Inactive*/+; Cher Active: w;*sns*-Gal4/+;UAS-*cher active*/+; Cher Active; TOR DN: w;*sns*-Gal4/+;UAS-*cher active*/UAS-*tor-dominant-negative*. One-way ANOVA plus Tukey’s multiple comparisons test: *: p < 0.05. **E** Combination of Cher Active with Hippo, which results in the inhibition of Yorkie (Yap/Taz) caused a significant reduction of nephrocyte size, even below the cell size of Cher Inactive nephrocytes. Cher Inactive: w;*sns*-Gal4/+;UAS-*cher Inactive*/+; Cher Active: w;*sns*-Gal4/+;UAS-*cher active*/+; Cher Active; HIPPO: w;*sns*-Gal4/+;UAS-*cher active*/UAS-*hippo(dMST)*.*flag*. One-way ANOVA plus Tukey’s multiple comparisons test: **: p < 0.01; ****: p < 0.0001. **F** FITC Albumin assays revealed a significant size decrease when dominant negative TOR is expressed exclusively. Control: w;*sns*-Gal4/+;UAS-*dicer2*/+; TOR DN: w;*sns*-Gal4/+;UAS-*dicer2*/UAS-*tor-dominant-negative*. Student’s t-test: ****: p < 0.0001. **G** Expression of Hippo also resulted in a significant size decrease as assessed with FITC Albumin assays. Control: w;*sns*-Gal4/+;UAS-*dicer2*/+; HIPPO: w;*sns*-Gal4/+;UAS-*dicer2*/UAS-*hippo(dMST)*.*flag*. Student’s t-test: ***: p < 0.001.

### Excessive increase of Cheerio and Filamin B levels resulted in a pathological effect

Cheerio and Filamin B exhibit a protective role during injury in nephrocytes. Previous studies have shown that gain-of-function mutations in the mechanosensor TRPC6 caused a severe podocyte phenotype (Reiser et al., 2005, p. 6; Winn et al., 2005, p. 6). To address whether an excessive increase of Cheerio activity also result in a pathological phenotype, we generated lines homozygous for UAS-Cheerio or UAS-Filamin B (i.e. with two copies), with the aim of increasing the activity of Cheerio/Filamin B. Expression of a double dose of Cheerio or Filamin B produced a pathological phenotype, as nephrocyte morphology was severely disrupted, FITC-Albumin uptake was significantly decreased and pupal development in the presence of AgNO_3_ was significantly delayed **(Figure 5B,C,D,E,F,G, Supp. Figure 5A,B,D,E)**. Interestingly, the previously observed hypertrophy phenotype is not present in the homozygous flies **(Figure 5A)**. Together with the data described above, this suggests that a moderate increase in Cheerio activation is beneficial and protective under conditions of injury, but beyond a certain level Cheerio activation becomes pathological. In further support of this, flies expressing the different Cheerio variants as single copy but raised at 28°C to increase protein levels, also presented with a severe morphological phenotype **(Supp. Figure 5C)**.

**Figure 5:**
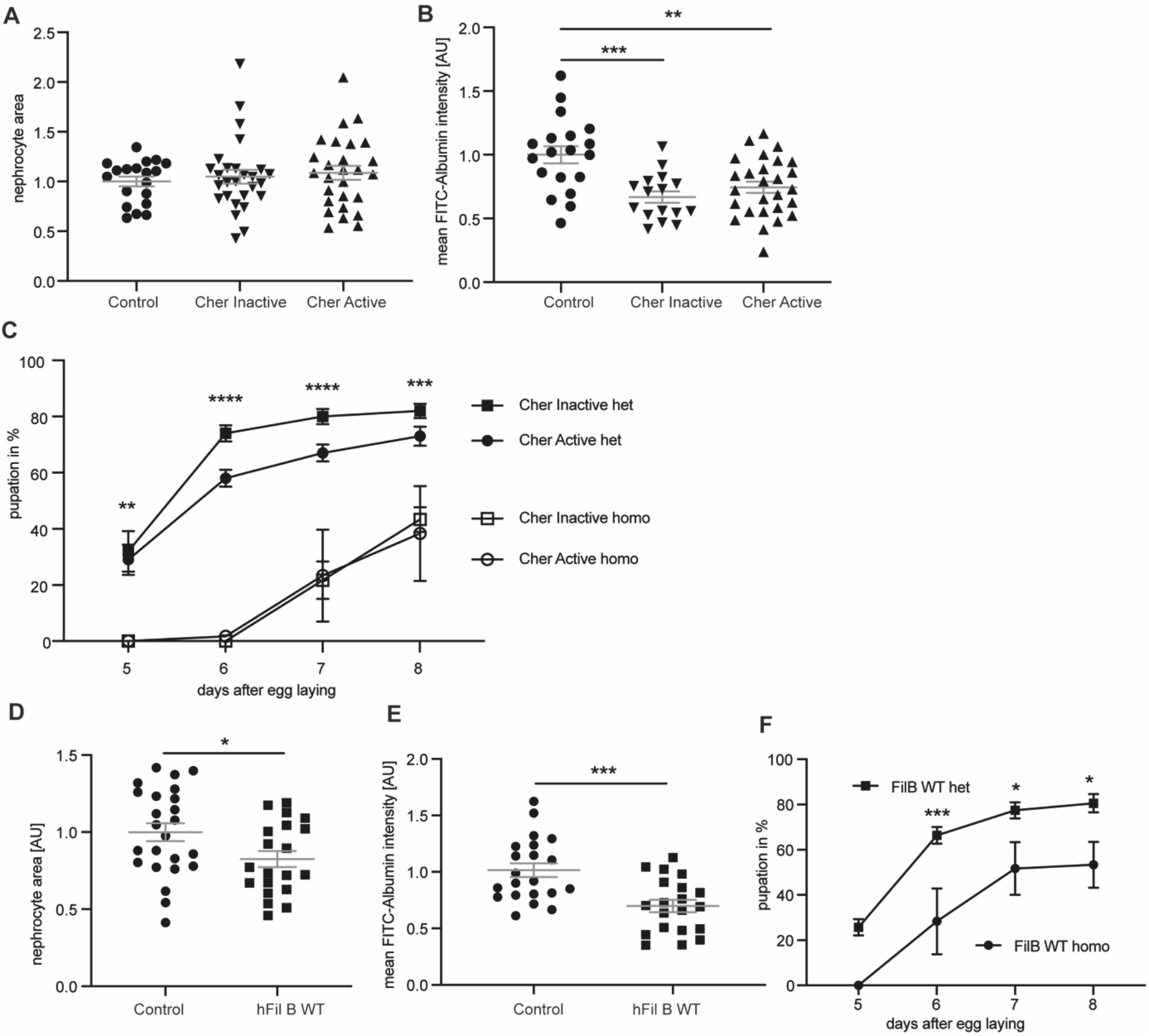
Excessive increase of Cheerio and Filamin B levels results in a pathological response. **A** Homozygous Active or Inactive Cheerio were used to increase the level of protein expression in nephrocytes. Comparison with control flies (w;*sns*-Gal4;MKRS/TM6B) revealed no difference in neprhocyte size. Cher Inactive: w;*sns*-Gal4;UAS-*cher-inactive;* Cher Active: w;*sns*-Gal4;UAS-*cher-active*. **B** FITC-Albumin uptake was significantly decreased in both genotypes (Inactive and Active Cheerio). One-way ANOVA plus Tukey’s multiple comparisons test: **: p < 0.01. **C** The AgNO_3_ toxin assay revealed a severely delayed pupation in homozygous flies when compared to their heterozygous controls. Cher Inactive het: w;*sns*-Gal4/+;UAS-*cher-inactive/*UAS-*dicer2;* Cher Active: w;*sns*-Gal4/+;UAS-*cher-active*/UAS-*dicer2;* Cher Inactive homo: w;*sns*-Gal4;UAS-*cher-inactive;* Cher Active homo: w;*sns*-Gal4;UAS-*cher-active*. Two-way ANOVA plus Tukey’s multiple comparisons test: **: p < 0.01; ***: p < 0.001; ****: p < 0.0001. **D** Homozygous expression of human Filamin B wildtype resulted in a significant size decrease. Student’s t-test: *: p < 0.05. Control: w;*sns*-Gal4;MKRS/TM6B; hFil B WT: w;*sns*-Gal4;UAS-*hFilamin B WT*. **E** FITC Albumin uptake was severely impaired by homozygous expression of human Filamin B wildtype. Student’s t-test: ***: p < 0.001. **F** Comparison of homozygous and heterozygous controls in the AgNO_3_ toxin assay also revealed severe filtration defects. Two-way ANOVA plus Tukey’s multiple comparisons test: *: p < 0.05; ***: p < 0.001.

**Supp. Figure 5:**
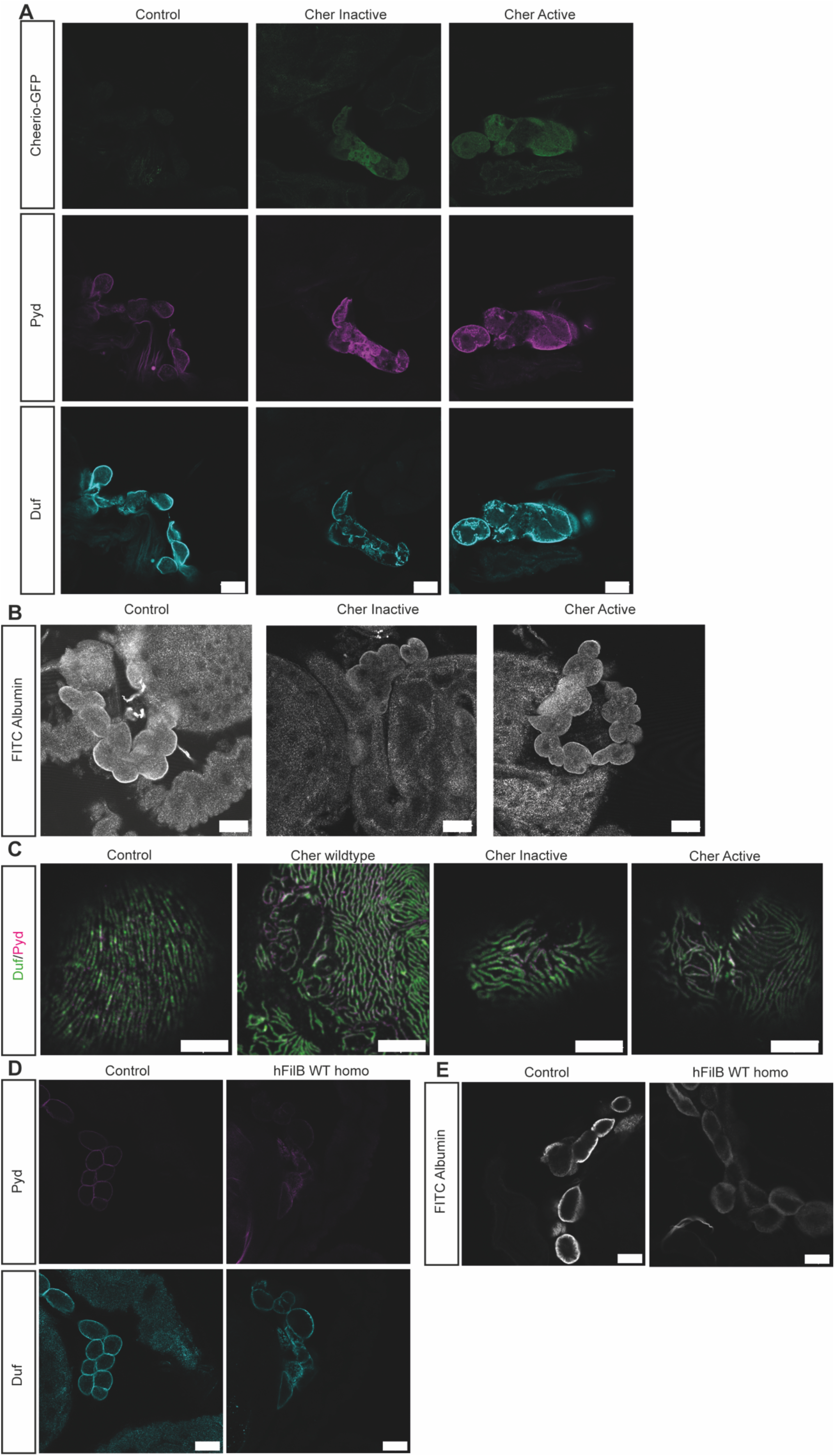
Excessive expression of Cheerio and human Filamin B result in a severe nephrocyte phenotype. **A** Immunofluorescence using a Duf and Pyd antibody revealed a morphological phenotype, when Cheerio Active and Inactive are homozygously expressed. Scale bar = 25 µm. Control: w;*sns*-Gal4;MKRS/TM6B; Cher Inactive: w;*sns*-Gal4;UAS-*cher-inactive;* Cher Active: w;*sns*-Gal4;UAS-*cher-active*. **B** FITC Albumin assays show a decreased uptake capacity in nephrocytes expressing either Cheerio active or Cheerio inactive. Scale bar = 25 µm. **C** Immunofluorescence staining using Duf and Pyd antibodies revealed morphological changes in nephrocytes expressing Cheerio wildtype, inactive and active at 28°C. Control: w;*sns*-Gal4;UAS-*dicer2*, Cher wildtype: w;*sns*-Gal4/UAS-*cher-wildtype-MSR*;UAS-*dicer2*/+, Cher Inactive: w;*sns*-Gal4/+;UAS-*dicer2*/UAS-*cher-inactive-MSR*, Cher Active: w;*sns*-Gal4/+;UAS-*dicer2*/UAS-*cher-active-MSR*. Scale bar = 5 µm. **D** Immunofluorescence staining with Duf and Pyd antibodies revealed changes of morphology in nephrocytes expressing human Filamin B wildtype. Scale bar = 25 µm. Control: w;*sns*-Gal4;MKRS/TM6B; hFil B WT: w;*sns*-Gal4;UAS-*hFilamin B WT*. **E** FITC Albumin uptake assays revealed a significant filtration defect after homozygous expression of human Filamin B wildtype. Scale bar = 25 µm.

## Discussion

Recent studies revealed an important role of the mechanosensor Filamin in podocyte biology, as both Filamin A and B were described to be upregulated under conditions of increased mechanical force, in several mammalian injury models and glomerular patient tissue (Greiten et al., 2021; Koehler et al., 2020; Okabe et al., 2021). In addition, it was previously shown that a nephrocyte-specific loss of Cheerio did not result in any change to nephrocyte diaphragm integrity or filtration function (Koehler et al., 2020). However, the function of Cheerio/Filamin in nephrocytes/podocytes is not known.

Based on these findings we asked whether Cheerio/Filamin exhibits a protective role during injury. To investigate this hypothesis, we utilized the *Drosophila* nephrocyte model and examined the functional role of Cheerio, the Filamin homologue in the fly. Our data show that an over-activation of the mechanosensor domain caused a hypertrophy phenotype in nephrocytes, but did not impact on nephrocyte diaphragm integrity or filtration function. The link between Filamin B and the hypertrophy phenotype was also recently observed in a mouse model investigating secondary injured podocytes (Okabe et al., 2021). In a novel partial podocytectomy mouse model (loss of a subpopulation of podocytes) Okabe et al. (May 2021) induced podocyte injury only in a subset of podocytes resulting in secondary damage in the remaining cells. Additional transcriptomics analysis revealed an upregulation of Filamin B and a hypertrophy phenotype in the remaining podocytes (Okabe et al., 2021).

An increased fluid flow causes an increased hydrostatic pressure in the capillaries during injury and disease. As a result, the filtration barrier, including the glomerular basement membrane, endothelial cells and podocytes will expand to reduce the filtrate flow per unit filtration area. However, this expansion is limited and will be followed by an increase of shear stress in the filtration slits and between podocytes in the Bowman’s capsule, resulting in podocyte detachment and loss (Butt et al., 2020; Kriz and Lemley, 2017). This loss of a podocyte subpopulation causes an additional increase of biomechanical forces and the remaining podocytes need to activate adaptive mechanisms to remain attached to the basement membrane and to cover the blank basement membrane areas (Koehler and Rinschen, 2021; Kriz and Lemley, 2017).

The upregulation of the mechanosensor Filamin B and the observed hypertrophy phenotype in podocytes and nephrocytes further support this hypothesis. Hypertrophy was previously described to serve as a protective mechanism during podocyte injury, as podocytes try to cover blank capillaries after loss of neighbouring cells by increasing their cell size (Puelles et al., 2019; Wiggins et al., 2005).

Interestingly, Cheerio translocates and accumulates at the cell cortex upon injury in nephrocytes. Previous studies in fibroblasts reported that active Filamin accumulates at the cell cortex, while inactive Filamin remains mainly cytoplasmic (Nakamura et al., 2014; Razinia et al., 2012). In line with that, we also found that active Cheerio localized to the nephrocyte cortex, whereas the inactive variant remained cytoplasmic **(Figure 6)**. Interestingly, inactive Cheerio (variant engineered to be less responsive to mechanical force) accumulates at the cortex in injured nephrocytes (upon loss of Duf and Sns). It is possible that during injury the forces experienced by Cheerio are such that even this ‘inactive’ variant is activated and translocates to the cell periphery and nephrocyte diaphragm.

**Figure 6:**
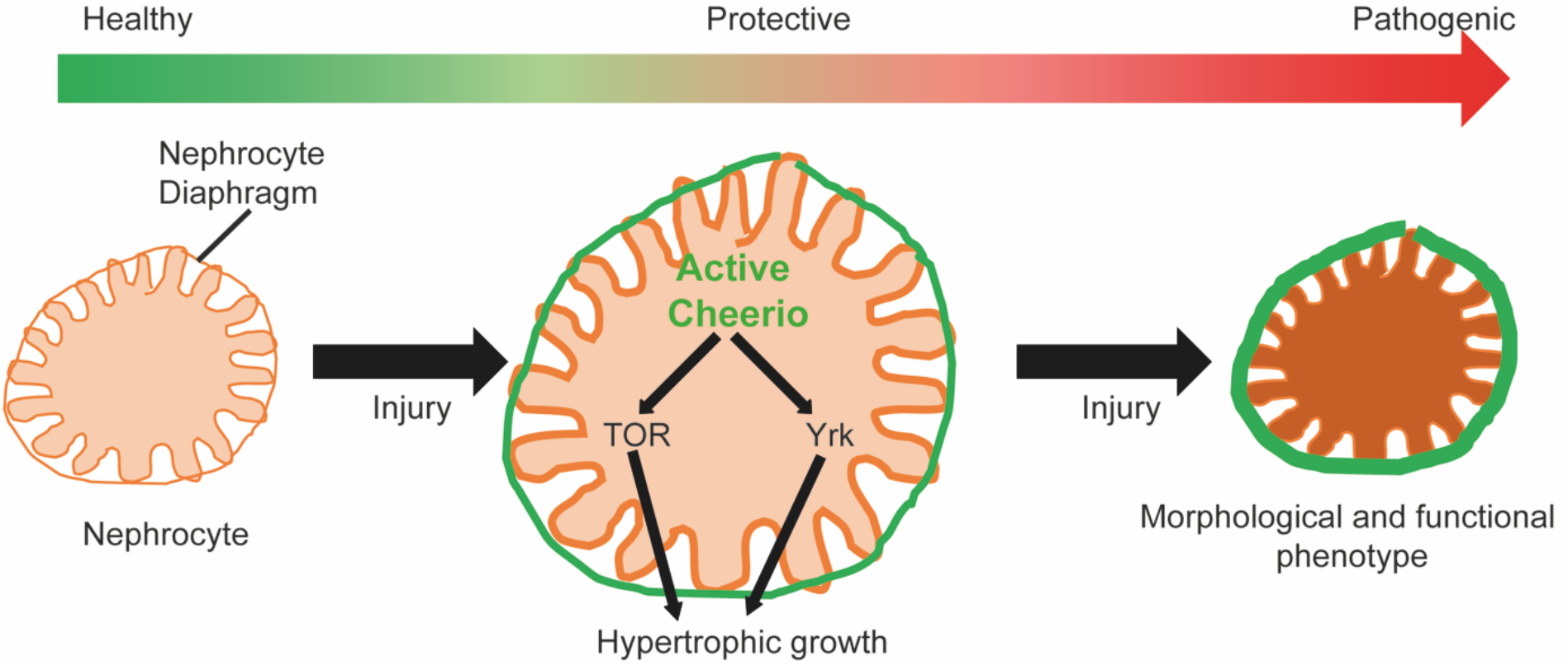
The mechano-protective role of Cheerio in *Drosophila* nephrocytes. Cheerio is active during injury and accumulates at the cell periphery. This activation and the increased protein levels result in a mechano-protective effect via hypertrophic growth, mediated via TOR and Hippo signalling. However, Cheerio levels need to be tightly controlled, as excessive increase of protein levels results in a shift from the protective to a pathogenic effect, including a morphological and functional phenotype.

Our data also shows that the expression of active Cheerio caused a rescue of nephrocyte function in Sns and Duf depleted cells **(Figure 6)**. Similar effects were observed when expressing the inactive variant. Of note, inactive Cheerio seems to be activated during injury based on its localisation at the periphery.

In addition, expressing human Filamin B suggested that this protective effect is evolutionarily conserved. As the human Filamin B construct also possessed the ACB and expression of human Filamin B rescued cell size during injury, we speculate that the ACB plays a role in cell size control.

Moreover, our data provides evidence that the levels of Cheerio and Filamin B need to be tightly controlled as excessive levels result in a severe nephrocyte phenotype, indicating a threshold where protective effects tip toward a pathological condition **(Figure 6)**. We have also provided evidence that hypertrophy might be mediated by TOR and/or Hippo pathways **(Figure 6)**. In line with our data, overactive mTOR caused a hypertrophy phenotype in mouse podocytes, which is associated with a protective effect during injury (Puelles et al., 2019). Moreover, several studies show that loss of YAP in podocytes causes a glomerular disease (FSGS) phenotype (Schwartzman et al., 2016), that YAP activation has a protective effect during injury (Meliambro et al., 2017) and that nuclear localization/activation of YAP has a pro-survival effect in podocytes (Bonse et al., 2018). Interestingly, in FSGS patient tissue, mTOR target genes were upregulated (Puelles et al., 2019), while inactive phospho-YAP levels were increased (Meliambro et al., 2017; Schwartzman et al., 2016). These findings, together with the upregulated Filamin B levels in podocyte models, indicate that the mechanisms investigated within this project might be highly evolutionarily conserved and that TOR and YAP might be downstream targets of Filamin in mammals as well.

In a recent study, Greiten et al. described the functional role of Filamin A in podocytes, and showed that loss of Filamin A caused a rearrangement of the actin-cytoskeleton and decreased expression levels of focal adhesion associated proteins (Greiten et al., 2021). They also showed increased Filamin A levels during hypertension in mouse models and hypertensive glomerular patient tissue (Greiten et al., 2021), again supporting our hypothesis of a protective role of Cheerio and Filamin during injury, when expressed at moderate levels.

Taken together, our data show that Cheerio is activated during injury, which result in a protective hypertrophy phenotype, possibly mediated via TOR and Hippo signalling **(Figure 6)**. However, Cheerio and Filamin levels need to be tightly controlled, as an excessive increase of activity result in a shift from the protective effect to a pathological response **(Figure 6)**.

## Author contribution

S.K. conceived the study, performed experiments, analysed the data, made the figures and drafted the paper, B.D. revised the study critically for important intellectual content, S.K. and B.D. revised the paper; both authors approved the final version of the manuscript.

## Acknowledgement

The Anti-Pyd monoclonal antibody developed by Fanning (Choi et al., 2011) was obtained from the Developmental Studies Hybridoma Bank, created by the NICHD of the NIH and maintained at The University of Iowa, Department of Biology, Iowa City, IA 52242.

